# A novel HLA Class II presentation prediction algorithm deciphers immunogenic CD4 epitopes specific to KRAS G12C

**DOI:** 10.1101/2024.12.06.627073

**Authors:** Daniel Sprague, Meghan G. Hart, Joshua Klein, Sonia Kounlavouth, Rahulsimham Vegesna, Melissa Rotunno, Lauren D. Kraemer-Tardif, Rita Zhou, Lindsey Kemp, Adrienne C. Greene, Joshua Araya, Alexis Mantilla, Bukola Adeoye, Calixto Dominguez, Andrew R. Ferguson, Melissa L. Johnson, Matthew J. Davis, Monica Lane, Christine D. Palmer, Karin Jooss, Ankur Dhanik

## Abstract

Accurate prediction of peptide presentation by HLA molecules is important for generation of effective individualized cancer vaccines and immunotherapies. While presentation prediction algorithms for HLA class I have been successfully applied in the context of such therapies, improved prediction algorithms for class II are needed. EDGE-II is a novel algorithm based on a protein large language model that has a learned allele deconvolution network trained on existing and new immunopeptidomics data. It delivers state-of-the-art performance on prediction of peptide presentation by HLA class II and immunogenicity elicited by CD4+ T-cell epitopes. In a patient with a KRAS G12C positive tumor treated with a KRAS G12C targeting immunotherapy, EDGE-II identified KRAS G12C class II neoantigens that elicited clonally expanded CD4^+^ T cells with cytotoxic transcriptional profiles post-vaccination. EDGE-II could play an important role in the development of effective cancer immunotherapies by elucidating an enriched understanding of the immunopeptidome.

## Introduction

Immunotherapy leverages T cells to target tumor neoantigens and offers a promising approach for the treatment of patients with cancer^1^. However, efficacy of the standard immune checkpoint blockade based immunotherapy is limited to the minority of patients across solid tumors and this associates with low levels of neoantigen-specific T cell infiltration into the tumor^1,2^. Recent advances in tumor neoantigen prediction^3^ and vaccine platforms^4^ has enabled the development of vaccine-based immunotherapies to drive neoantigen-specific T cells in patients with cancer. Individualized cancer vaccines, targeting multiple mutations unique to a particular patient is an approach increasingly being explored in clinic. Alternatively, an “off-the-shelf” immunotherapy that targets shared neoantigens arising from common single-nucleotide variants, such as KRAS mutations, is a highly desirable treatment approach for patients with KRAS mutation-positive solid tumors^5^. While studies indicate that cytotoxic CD8^+^ T cells directly targeting tumors presenting neoantigens are critical to tumor control and clearance^6^, the role of CD4^+^ T cells is still not fully understood^7^. In a melanoma study with adoptive T cell therapy, tumor regression was observed in a patient treated predominantly with CD4^+^ T cells^7^. Previous work has shown that CD4^+^ T cells can either directly eradicate tumor cells expressing HLA class II through cytolytic activity or indirectly by infiltrating the tumor invasive margins and interacting with HLA class II expressing professional antigen presenting cells (APCs), promoting an interferon-gamma (IFNγ)-induced anti-tumor response^8,9^. Overall, CD4^+^ T cell activation is believed to be a key component necessary for driving a broad and durable anti-tumor response^10^. The ability to identify putative HLA class II restricted tumor-specific T cell epitopes is therefore highly desirable to extend the design of potent cancer immunotherapies to target both class I and II epitopes.

Several algorithms exist that predict peptide presentation by HLA Class I with high positive predictive value (PPV) and have been used in a clinical setting^3,11–14^. For example, EDGE^TM^ was used in the development of individualized cancer vaccines that have demonstrated preliminary evidence of therapeutic benefit to patients with advanced cancer^4^. However, predicting peptide presentation by HLA class II has proven more challenging due to limited immunopeptidomics data (used for training data in predictive model construction) and less constrained peptide-HLA binding principles than for class I^11,15,16^. Several algorithms have made progress over recent years^16–21^. Here, we present a new algorithm called EDGE-II for predicting peptide presentation by HLA class II. EDGE-II leverages transfer learning from a pre-trained protein large language model and conditions its predictions on full sample genotypes using learned multi-allele deconvolution. We demonstrate that EDGE-II achieves a state-of-the-art PPV and strong association with immunogenicity as compared to publicly available algorithms.

Recently, several KRAS G12 mutants were shown to elicit CD4^+^ T cell responses in patients with cancer^22^ and such mutations, especially G12C, are common^23,24^. This provides an opportunity for targeting class II presented mutant peptides with immunotherapies such as vaccines^22,25–27^. In our clinical study^28^ (<u>NCT03953235</u>), we observed KRAS G12C-specific class II-restricted CD4^+^ T cell responses in a patient treated with a KRAS G12C targeting vaccine. EDGE-II predicted presentation of G12C mutant peptides by HLA Class II alleles and identified a motif that is important for peptide presentation. The DQ class II alleles in the patient’s HLA genotype were associated with the strongest presentation probability for KRAS G12C mutant peptides.

In this work, we describe EDGE-II and show performance improvements over best public algorithms (available to us) on both immunopeptidomics and immunogenicity datasets. Using KRAS G12C as an example, we show that EDGE-II offers an opportunity to generate neoantigen-directed immunotherapies capable of driving broad and robust tumor-specific CD4^+^ T cells in addition to CD8+ T cell responses.

## Results

### EDGE-II delivered state-of-the-art HLA class II presentation predictions

Recent advancements in machine learning have enabled development and use of pre-trained protein large language models (PLLM) for a variety of biological inference tasks because these models embed the complex physicochemical and structural information of protein sequences^29,30^. Fine-tuning the parameters of a PLLM for inference tasks in protein biology, rather than starting from random initialization, generally increases performance of predictive models^31^. BERTMHC^16^ is a HLA class II presentation model that utilizes a prior PLLM, TAPE^32^, and demonstrated improved predictions over NetMHCIIpan4.0^11^. We sought to build a sequence-only class II algorithm, named EDGE-II, using the ESM2 PLLM^33^ (**Fig. 1A**). To test if ESM2 is more effective for class II presentation prediction than the TAPE model used in BERTMHC, we built a model using either TAPE (38M parameters, 512 hidden dimensions) or the closest version of ESM2 (35M parameters, 480 hidden dimensions), and trained and evaluated on the well-known Reynisson *et al.* class II presentation data^11,34^ (**Fig. 1B**). The predictive model trained on ESM2 significantly outperformed TAPE (**Fig. 1C**).

**Figure 1:**
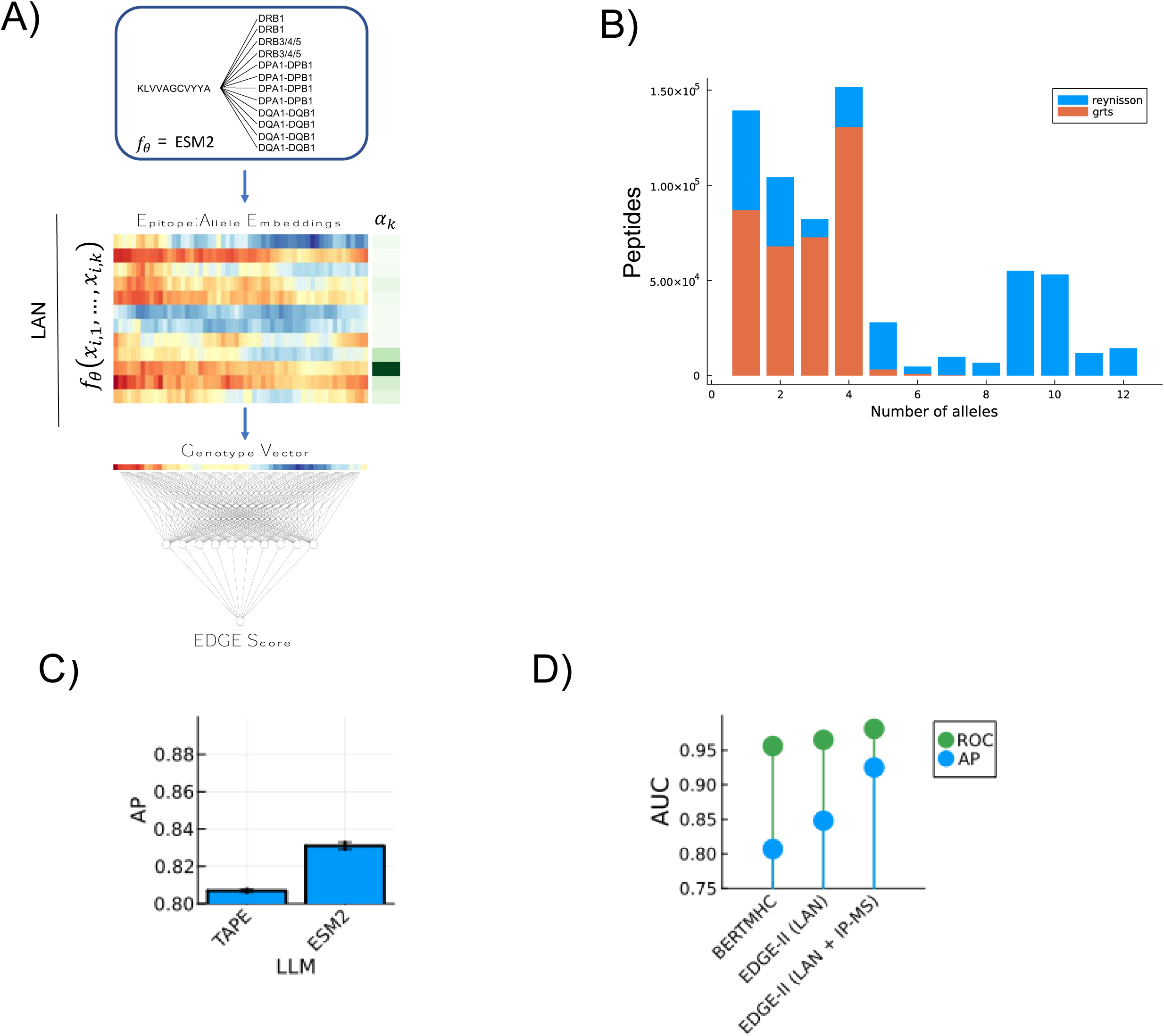
EDGE-II uses a protein language model and learned multi-allele deconvolution via Learned Allele Network (LAN). **A)** EDGE-II leverages a state-of-the-art PLLM and a novel learned allele network (LAN) that delivers genotype specific predictions for a given epitope. Left: Model architecture, where a peptide sequence is independently concatenated with the pseudosequences of all HLA class II alleles present in a genotype and passed through the ESM2 PLLM. EDGE-II’s deconvolution strategy is unique in that it is learned and operates on the PLLM embeddings rather than on predicted scores (middle). *a^k^* are the allele weights emitted by the learned allele weight network and are unique to a given genotype. **B)** Dataset size parsed by number of alleles in each sample. Data collected from Reynisson *et al.* and data generated inhouse is shown. **C)** Average precision for classifiers trained using TAPE vs ESM2 on the validation split of the Reynisson *et al*. data. Error bars represent bootstrap samples. **D)** Validation metrics of EDGE-II on the hold-out split of the Reynisson *et al*. data compared against the reported values in BERTMHC.

Following the inclusion of ESM2, we sought to improve the deconvolution of multi-allelic immunopeptidomics data typically derived from HLA genotyped human tissues or cell lines. Multi-allelic (MA) data, in addition to the single-allelic (SA) data that are typically derived from monoallelic cell lines, have become a key component of training HLA presentation algorithms for class I and class II. However, this MA data require a mathematical method to produce a single presentation probability across the multiple alleles. Within the context of HLA presentation, we refer to such methods as allele deconvolution. Prior class II methods enforced heuristics such as iterative pseudo-labeling^34^ or per-allele maximal score deconvolution^16^. We sought to determine if EDGE-II could learn to deconvolute multiallelic data based off the alleles present in the sample and the putative peptide being considered. To do this, we implemented a Learned Allele Network (LAN) that operates on the latent epitope:allele embeddings from ESM2 rather than on the final output of prediction probabilities, as in BERTMHC, NetMHCIIpan4.3, and existing class I models (**Fig. 1A**, Methods). In the LAN, the allele weights, *a_k_*, are dependent on the complete genotype present in a sample and are calculated at inference time by incorporating a specialized auxiliary attention network into EDGE-II. Since standard MA data are generated using pan-specific antibodies, we next leveraged immunoprecipitation (IP) with HLA-DR/DP/DQ-specific antibodies to generate mass spectrometry (MS) presentation data to not only increase the size of the training dataset, but also the resolution of the peptide-HLA associations in it (**Fig. 1B**).

We first trained and evaluated EDGE-II on the Reynisson *et al.* dataset used to train both NetMHCIIpan4.1 and BERTMHC. With the LAN’s inclusion in EDGE-II, we achieve an Average Precision (AP) = 0.848 ± 0.001, ROC-AUC = 0.963 ± 0.003 which is a substantial gain in performance over the maximal output probability deconvolution approach in BERTMHC (AP = 0.807 ± 0.001, ROC-AUC = .954 ± 0.0004) or EDGE-II with maximal output probability deconvolution instead of LAN (AP = 0.831 ± 0.002, ROC-AUC = 0.958 ± .0003; **Fig. 1D**). Inclusion of the HLA-DR/DP/DQ-specific immunopeptidomics data as an intermediate training step between SA training and MA deconvolution training led to substantial improvements in ROC-AUC and AP compared to the same model architecture that did not include these data, with EDGE-II achieving a final validation AP = 0.925 ± 0.005 and ROC-AUC = 0.981 ± 0.004 on the Reynisson validation data (**Fig. 1D**). Breaking performance down to individual alleles revealed substantial improvements in EDGE-II as compared to BERTMHC across most alleles (**Fig. 1D)**.

We next applied EDGE-II to the independent test set of mass spectrometry data curated by Cheng *et al.* when evaluating BERTMHC (**Fig. 2A**). We found that EDGE-II (AP = 0.684 ± 0.007, ROC-AUC = 0.898 ± 0.004), trained on the same dataset as BERTMHC and excluding our IP-MS data, substantially outperformed BERTMHC (AP = 0.648 ± 0.006, ROC-AUC = 0.889 ± 0.002), NetMHCIIpan4.3 (AP = 0.577 ± 0.002, ROC-AUC = 0.821 ± 0.003), and MixMHC2pred-2.0 (AP = 0.442 ± 0.003, ROC-AUC = 0.790 ± 0.003). With the inclusion of the IP-MS data while training, EDGE-II achieved a test AP of 0.775 ± 0.004 and ROC-AUC of 0.934 ± 0.002 (**Fig. 2B**). Per-allele performance comparisons show substantial improvements in the worst performing alleles from MixMHC2pred-2.0 and NetMHCIIpan4.3 (**Fig. 2C**).

**Figure 2:**
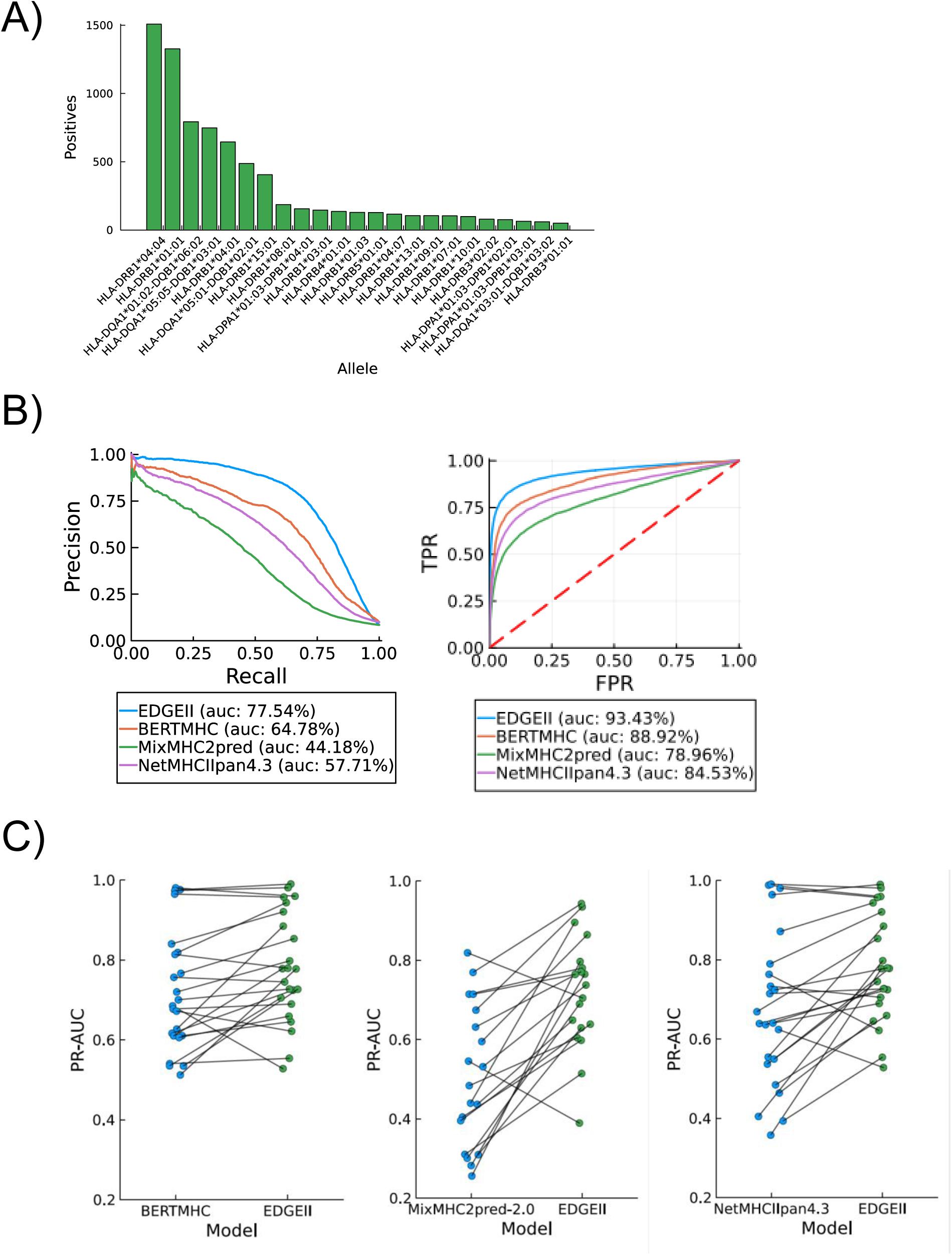
EDGE-II has improved predictive performance over available published algorithms. **A)** Distribution of peptides across HLA class II alleles in the independent test set curated by Cheng *et al*. **B)** Precision-recall and ROC curves on the IEDB sourced independent test set curated by Cheng *et al.* for evaluation of BERTMHC. NetMHCIIpan4.3, and MixMHC2pred-2.0 also included for this analysis. **C)** Per allele PR-AUC between EDGE-II and BERTMHC (left), MixMHC2pred-2.0 (middle), and NetMHCIIpan4.3 (right).

### EDGE-II predicted immunogenic neoepitopes presented by HLA class II

We next sought to determine if EDGE-II scores based only on sequence data are predictive of immunogenicity for class II presented epitopes. Incorporation of additional covariates such as gene expression and cleavage signatures generally improve prediction performance on immunopeptidomics data for HLA class I. However, in the case of HLA class II, natural peptide processing is decoupled from the eventual presentation of peptide by HLA class II on professional APCs. Several HLA class II predictors exclude common class I covariates other than the peptide, HLA alleles, and optionally the peptide context^19,20^. Gene expression is a common covariate used in HLA class I predictors; however, a non-trivial proportion of class II restricted peptides lack detectable RNA expression, of which a significant amount are extracellular proteins^17^. Last, limiting the number of conditioning predictors makes the algorithm more widely applicable.

To assess association between EDGE-II score and CD4^+^ immunogenicity in an individualized cancer vaccine context, we gathered ELISpot response data with full HLA class II genotypes from Ott *et al.*^35^ to determine if EDGE-II scores are positively associated with immunogenicity (**Fig. 3A**). Ott and colleagues generated individualized mutanome targeted peptide vaccines with the goal of inducing CD4^+^ and CD8^+^ T cell responses in melanoma patients, from which ELISpot T cell response data were collected. We therefore sought to evaluate the ability of EDGE-II to predict the induction of CD4^+^ T cell responses post vaccination. For each peptide in the ELISpot response data, we took a sliding 15 amino acid (aa) window across the peptide and calculated an EDGE-II score. The maximum score across the windows was then saved as the final presentation prediction for that peptide.

**Figure 3.**
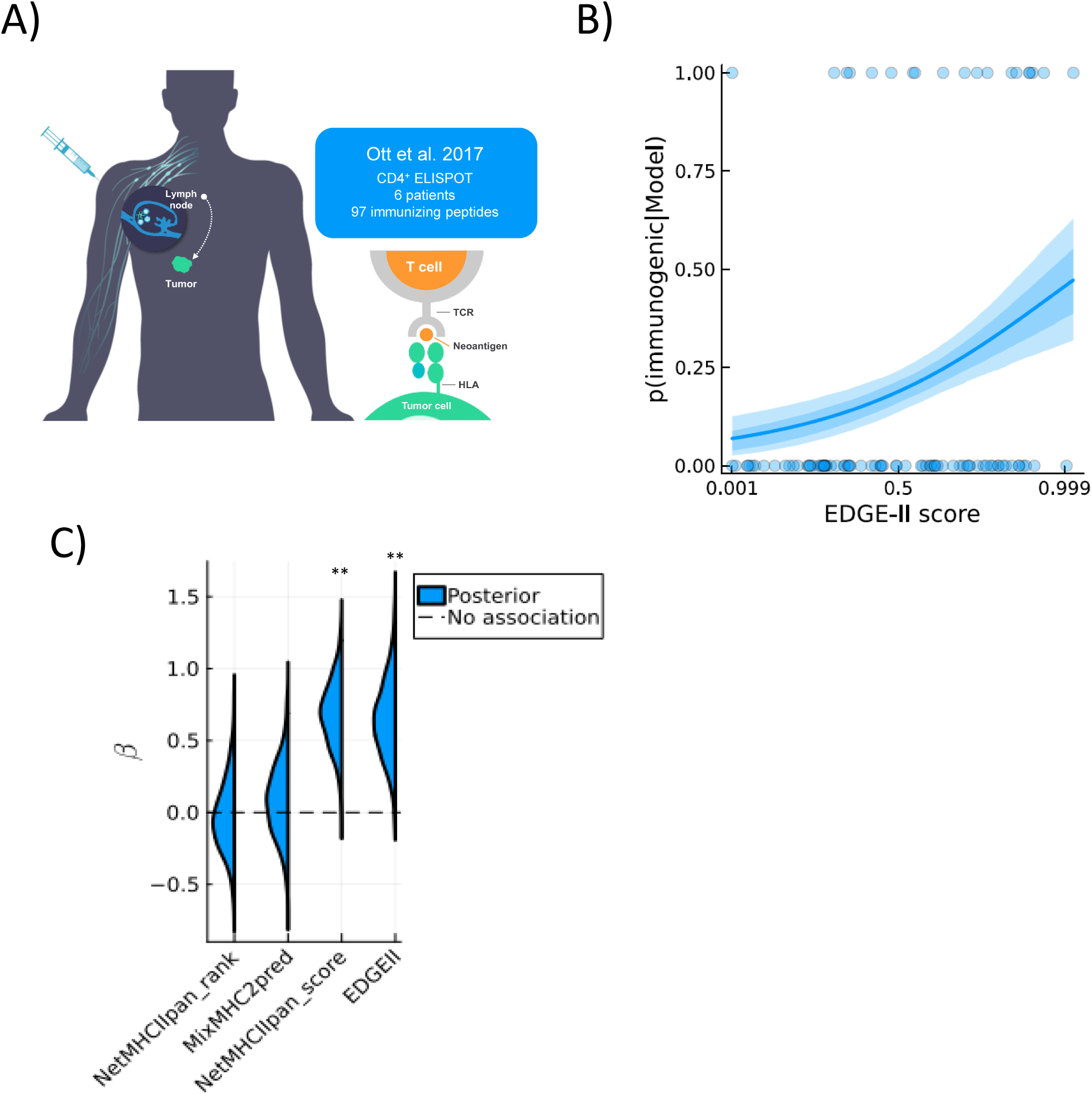
EDGE-II discriminated immunogenic and non-immunogenic neoepitopes. **A)** Description of the Ott et al. dataset used for immunogenicity analysis. **(b)** Posterior predictive distribution from a Bayesian explanatory regression of EDGE-II scores against functional ELISPOT data. Dots correspond to individual peptides that were ELISPOT positive (y = 1) or negative (y = 0). Dark blue represents 50% CI and light blue 90% CI. **C)** Sampled posterior distributions for EDGE-II and other algorithms from the Bayesian logistic regression analysis. *β* is the regression coefficient associated with model score. Predictors input into each explanatory regression were standardized such that coefficients are comparable to standard unit change.

EDGE-II scores were positively associated with immunogenicity over the full dataset. EDGE-II gave 9/97 peptides a presentation probability greater than 0.99, for which the expected probability of CD4 immunogenicity is 0.41 ± 0.12 90%CI (**Fig. 3B**). In contrast, the lowest scoring 10% of peptides in the dataset have a predicted probability of immunogenicity of 0.05 ± 0.04 90%CI (**Fig. 3B**). The percentile rank-based scores from MixMHC2pred-2.0 and NetMHCIIpan4.3 had almost no association with immunogenicity in this dataset, however presentation probabilities from NetMHCIIpan4.3 had significant and positive association that was comparable to EDGE-II (**Fig. 3C**). On these limited but high quality CD4 immunogenicity data, EDGE-II demonstrated significant association with functional T cell activity, despite using only sequence features. Furthermore, this analysis revealed that low scoring EDGE-II peptides have low likelihood of generating a CD4^+^ T cell response, whereas the highest scoring peptides are on average 8 times more likely to generate a CD4^+^ T cell response.

### Case Study: EDGE-II deciphered CD4 immunogenicity driven by KRAS G12C epitope

Driver mutations in the KRAS gene are prevalent in cancer and targeting KRAS G12C mutations with small molecule inhibitors has provided clear therapeutic benefit to patients^36,37^. However, acquired resistance can be observed within months of treatment initiation and pose a challenge for providing long-term clinical benefit to patients with cancer^38^. An “off-the-shelf” cancer vaccine targeting KRAS mutations could potentially offer an alternative or additive therapeutic approach for providing more durable benefit to these patients^39,40^. In a Phase 1/2 study, a patient with KRAS-G12C-positive NSCLC (S2) was treated with a KRAS G12C targeting immunotherapy. Consisting of a chimpanzee adenovirus vector and self-amplifying mRNA (samRNA) in combination with intravenous nivolumab and subcutaneous ipilimumab (**Fig. 4A**; <u>NCT03953235</u>) and had a 75% decrease in the KRAS G12C variant in circulating tumor DNA relative to baseline^28^.

**Figure 4.**
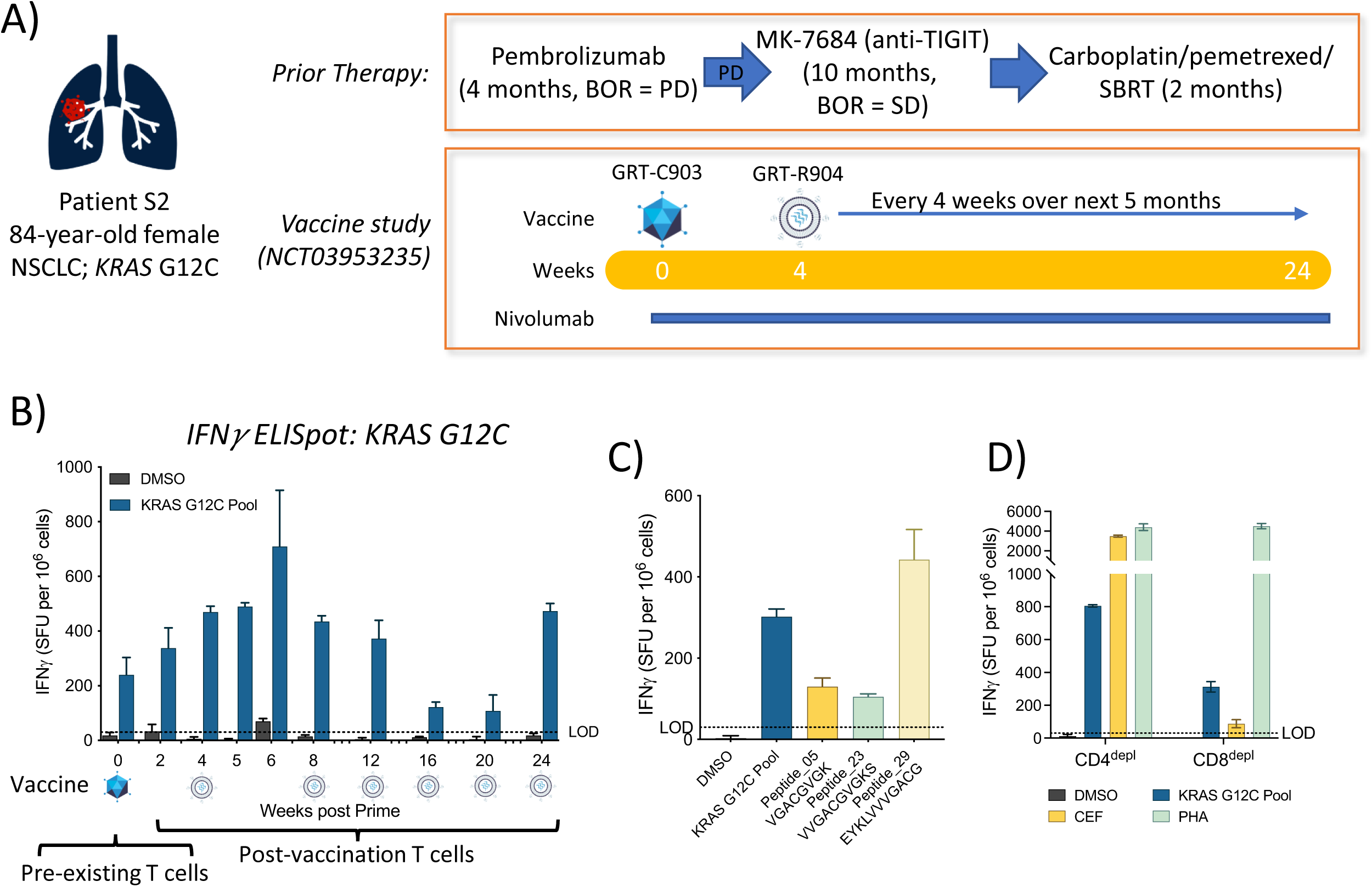
G12C-specific T cell responses were observed in NSCLC patient S2 treated with KRAS G12C targeting immunotherapy. **A)** Schematic showing patient S2 tumor profile and therapies prior to enrollment and vaccination in study NCT03953235^28^ B) T cell responses to KRAS G12C peptide pool assessed by *ex vivo* IFNγ ELISpot. Bar graphs (mean ± SD of replicate wells) show SFU/10^6^ cells for timepoints prior to (baseline; week 0), and after vaccination (weeks 2, 4, 5, 6, 8, 12, 16, 20, and 24). Assay LOD (30 SFU/10^6^ cells) is indicated by dotted line. Vaccination conditions are indicated by adenovirus and self-amplifying mRNA (samRNA) symbols. C) & D) T cell responses to KRAS G12C peptide pool and individual peptides or controls as assessed by *ex vivo* IFNγ ELISpot for PBMCs (C), CD4^depl^ PBMCs, and CD8^depl^ PBMCs (D) are shown. Bar graphs (mean ± SD of replicate wells) indicate SFU/10^6^ cells at post-vaccination timepoints. Assay LOD (30 SFU/10^6^ cells) is indicated by dotted line.

We initially assessed patient PBMC samples from available post-vaccination timepoints for putative KRAS G12C-specific CD8^+^ T cell responses using a pool of short (8-11aa) peptides as part of standardized immune monitoring (**Fig. 4B**; <u>NCT03953235</u>)^28^. Deconvolution of patient T cell responses to single 8-11mer peptides confirmed responses to an 8mer (Peptide_05), a 9mer (Peptide_23), and an 11mer (Peptide_29) with a nested core sequence (VGACGVGK) present in all 3 responsive peptides (**Fig. 4C**). Depletion of either CD4^+^ or CD8^+^ T cells from patient PBMCs revealed a mixed CD4^+^ and CD8^+^ T cell response to KRAS G12C 8mer-11mer peptide pools (**Fig. 4D**). Limited sample availability for patient S2 precluded further analyses using PBMC samples.

In addition to our observation, KRAS G12 mutants, including KRAS G12C, have recently been demonstrated to generate a class II-dependent immune response^22^. Despite these observations, we have observed that HLA class II models typically yield low presentation scores for KRAS G12C epitopes across alleles. NetMHCIIpan4.3, for example, predicts no strong binders, one weak epitope in DR, two weak epitopes in DP, and no DQ epitopes for the G12C mutation amongst MS supported alleles. We used EDGE-II to evaluate each allele in patient S2 to determine the likelihood of KRAS G12C peptide presentation by HLA class II. We found that EDGE-II yielded a high presentation score for KRAS G12C epitopes and particularly for HLA-DQA1*05:05-DQB1*03:01, with a score of 0.54 falling into the 99^th^ percentile for that allele (**Fig. 5A**). The maximally scoring peptide contained the nested core sequence (VGACGVGK) present in all responsive peptides described above (**Fig. 5B**). In general, we found DQ alleles to be more strongly associated with high EDGE-II scores as compared to DP and DR alleles (**Fig. 5C**).

**Figure 5:**
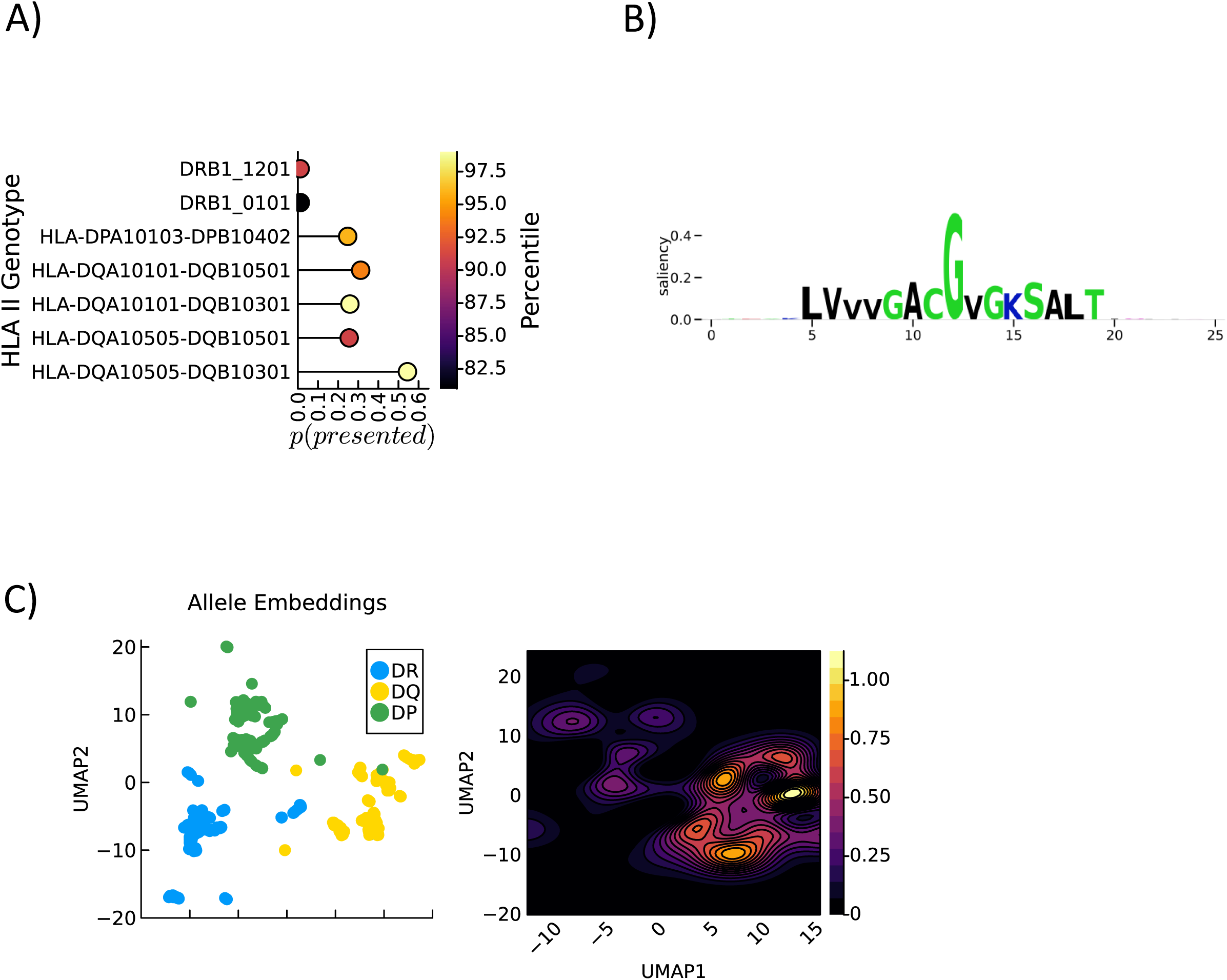
EDGE-II predicted HLA Class II alleles and peptide motif comprising KRAS G12C Class II neoantigens. **A)** Maximal presentation probability among all 15-mers containing the G12C mutation against the HLA genotype from a patient that received a vaccine containing KRAS G12C antigen. Presentation probability was scored per-allele. **B)** Saliency map of the first 26 amino acids of KRAS G12C computed by extracting the gradient of each 15-mer among G12C containing epitopes, over Patient S2’s HLA class II genotype (Methods). Larger saliency indicates greater importance to prediction of HLA class II presentation. **C)** Left: UMAP representation of all unique HLA alleles for which pseudosequences are available, colored by gene family. HLA alleles were embedded using pre-trained ESM2 (not EDGE-II). Right: Contour plots show EDGE-II scores as a function of the embedded HLA sequence-space, modeled as a 2D Gaussian Process (Methods).

TCR clonotype and transcriptional profiles were analyzed via single cell sequencing across multiple samples from patient S2 either stimulated with KRAS G12C pools or CEF (CD8^+^ T cell-specific responses to short Cytomegalovirus, Epstein-Barr virus and Influenza peptides) controls (**Supplementary Figure S1**). Cell line-based functional assays assessing function of the 95 most frequent TCR clonotypes via recombinant TCR (rTCR) expression were performed. Jurkat cells expressing patient S2 rTCRs co-cultured with K562 cells expressing each of the 6 patient-matched single class I alleles did not result in any positive responses to KRAS G12C 8mer-11mer peptide pools (data not shown). However, co-culture of rTCR-expressing Jurkat cells with class II matched KAS116 cells identified two TCR clonotypes (TCR969 and TCR995) that elicited increased expression of T-cell activation markers CD69 and CD25 on rTCR Jurkat cells in response to KRAS G12C peptide pools (both 8-11mer and 10-20mer pools) and KRAS G12C single peptides in 2 independent experiments (**Fig. 6A**). Importantly, rTCR functional assays confirmed responses to individual peptides (Peptide_05, Peptide_23, Peptide_29) observed in patient S2 via ELISpot (**Fig. 6A**).

**Figure 6:**
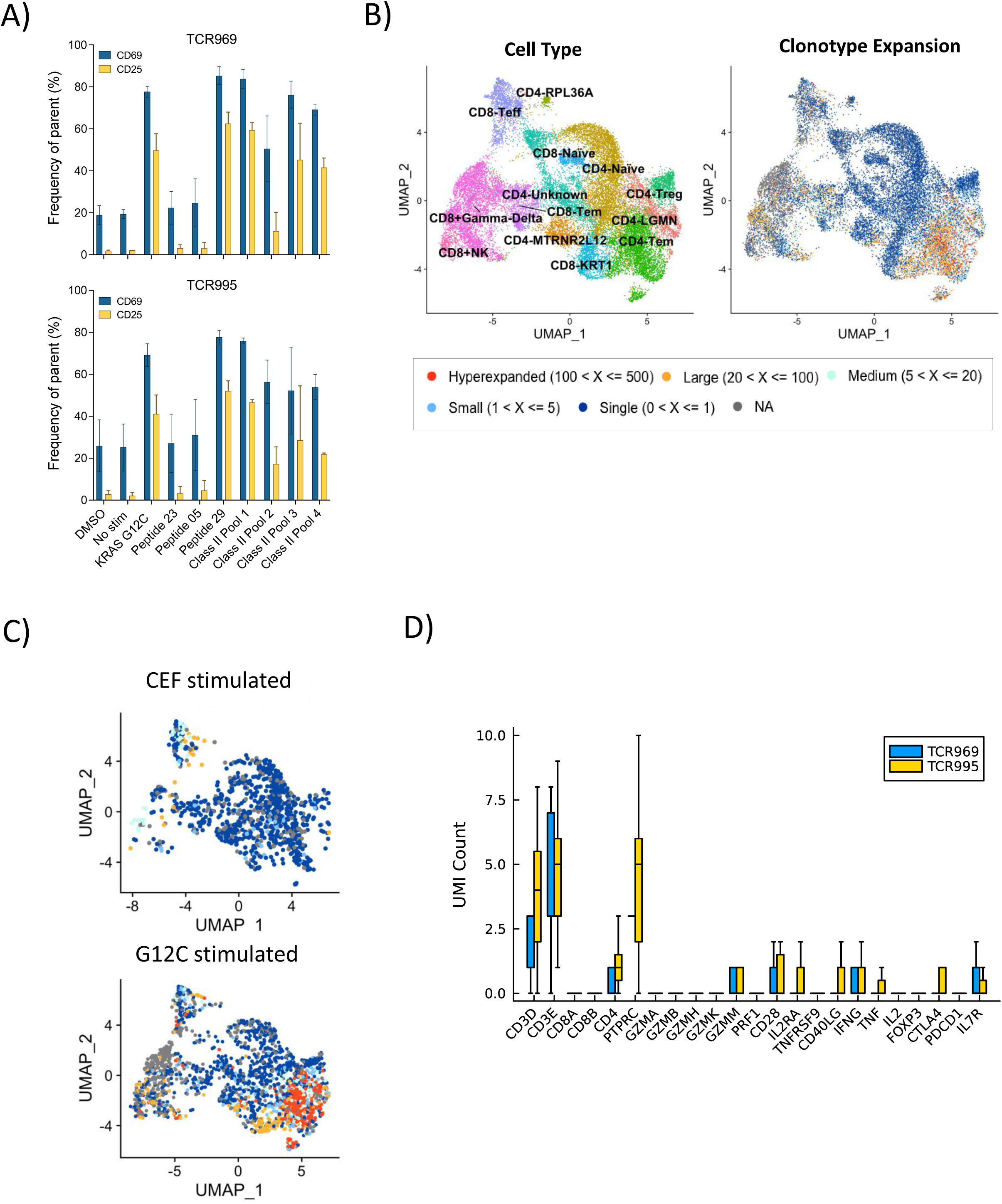
Expanded KRAS G12C specific CD4^+^ T-cells were observed in NSCLC patient S2 post-immunotherapy. **A)** Activation of Jurkat cells transduced with recombinant TCRs (rTCRs) from patient S2 co-cultured with antigen presenting cells pulsed with KRAS G12C pools and peptides or controls are shown. Bar graphs (median of replicate experiments ± SEM) show surface expression of activation markers CD25 (yellow) and CD69 (blue) as percent (%) positive populations. **B)** T cells from timepoint Dose-4 Day-1 visualized in two dimensions using UMAP and colored by clonal expansion. Top: Stimulation with CEF peptides. Bottom: Stimulation with G12C peptides. **C)** Approximately 23K T cells from multiple experimental conditions were visualized in two dimensions using UMAP (Methods). Left: annotation of different cell types from transcription profiles. Right: Same visualization with clonal abundance as the colormap. The T cells colored in red represent the clonotypes that are hyperexpanded (each unique clonotype is represented by 100 to 500 cells) and the T cells colored in blue represent the clonotypes that are singletons (each unique clonotype is represented by a single cell). **D)** UMI counts (expression level) of genes (x-axis) associated with T cell types and functional markers from single cell sequencing data. TCR969 (blue, *n*= 5 cells) and TCR995 (yellow, *n*= 7 cells) were experimentally validated for binding to a G12C peptide.

Transcriptional clustering analyses revealed distinct CD4^+^ and CD8^+^ T cell phenotype clusters with varying degrees of TCR clonotype expansion across all samples (**Fig. 6B**). As expected, in control samples stimulated with viral epitopes (CEF), proliferating clonotypes predominantly mapped to a population with CD8^+^ effector memory transcriptional profiles (**Fig. 6C**). In contrast, in patient PBMC samples stimulated with G12C peptides, hyper-expanding clonotypes predominantly mapped to CD4^+^ populations with effector memory transcriptional profiles, although expanding cells also clustered with CD8^+^ T cell populations, in line with a mixed CD4^+^/CD8^+^ G12C-specific T cell response observed in PBMCs (**Fig. 6C**). Importantly, the two functional TCRs identified during co-culture with class II matched cell lines (TCR969, TRC995) mapped to cell populations with a CD4^+^ cytotoxic transcriptional profiles (**Fig. 6D**), suggesting the presence of G12C-specific cytotoxic CD4^+^ T cell responses in patient PBMCs.

## Discussion

Accurate prediction of HLA class II epitopes is important for the development of therapeutic cancer immunotherapies that can induce potent and broad CD4^+^ and CD8^+^ anti-tumor T cell responses. HLA class II presentation has been shown to be positively associated with anti-tumor immune responses following cancer immunotherapy^35,41^. However, the prediction of peptides presented by HLA class II remains a challenging task due to the complexity of the underlying biology and relative scarcity of experimental immunopeptidomics data.

EDGE-II employs recent advances in the field of machine learning, particularly the large protein language model ESM2, a novel learned allele network (“LAN”) based deconvolution strategy, and DR/DP/DQ-gene specific immunopeptidomics training data to improve the state of the art in class II presentation prediction. Since EDGE-II is a sequence-only model, it does not depend on covariates such as gene expression, cleavage scores, etc. and therefore is widely applicable. Unlike other published models, the LAN provides EDGE-II the ability to deconvolute peptide presentation in the context of a patient’s full HLA genotype which results in improved model performance on multi-allelic (MA) data. While precise prediction of presented epitopes is important, the desired outcome of a shared or individualized cancer vaccine is a functional T cell response that recognizes and kills tumor cells. When applied to the Ott *et al.* dataset, for which full class II HLA genotypes are available, we found that EDGE-II was positively and significantly associated with immunogenicity. Interestingly, we found that rank-based scoring methodologies demonstrated no association with immunogenicity in this small but high-quality dataset. NetMHCIIpan4.3, which provides both presentation probability and percentile rank, shows association with presentation probability but not percentile rank. This suggests that the rank of a peptide for a given allele is less important than the overall estimate of presentation probability.

A recent study administering an individualized cancer vaccine in combination with immune checkpoint blockage therapy and chemotherapy in patients with NSCLC reported KRAS G12C-specific CD4^+^ T cell responses^42^. While no specifics on class II presentation were given, there was evidence of cytotoxic function based on CD107a and IFNg^+^ expression in one patient although there was no data evaluating direct killing. In agreement with these data, we observed expansion of KRAS G12C-specific CD4^+^ T cells with cytotoxic function in a patient with KRAS G12C-positive NCSLC post vaccination with a shared neoantigen immunotherapy regimen. Predicting presentation of KRAS G12C peptides by HLA class II has remained a challenge for prediction models. EDGE-II, however, predicted a strong HLA class II presentation signal for KRAS G12C peptides.

Evidence for direct or indirect anti-tumor activity conferred by CD4^+^ T cells^43–45^ targeting MHC Class II-positive and -negative tumors^46,47^ suggests that most potent vaccination strategies will benefit from induction of tumor-specific CD4^+^ and CD8^+^ T cell responses^10^. EDGE-II enables improved prediction of peptides presented by HLA class II and, thus, the rational development of cancer vaccines designed to induce broad T cell responses. Although complex and challenging, accurate prediction of the immunogenicity of tumor antigens is highly desirable for vaccine design. EDGE-II demonstrates not only an improved capability of predicting peptide presentation, but also presents a step forward towards predicting CD4 immunogenicity, with the overall goal of generating potent neoantigen-specific cancer vaccines and other immunotherapies that provide durable benefit to patients with cancer.

## Methods

### HLA class II presentation data

Data were collected from internal and publicly available sources to train EDGE-II. To validate the EDGE-II architecture against prior methods, we used previously published eluted ligand data without alteration^11,16^. This data contains 41 HLA class II alleles with peptides ranging from 13aa-21aa. This dataset sampled negative (not presented) epitopes randomly from the proteome at a prevalence of 10 negative examples for every presented epitope, for each allele or set of alleles, from the UNIPROT database. The allelic specificity of the data ranges from unambiguous single allelic data to full class II HLA genotypes containing 12 possible presenting alleles. Internal HLA class II presentation data were generated using HLA class II allele family specific immunoaffinity capture followed by mass spectrometry. All epitopes were therefore restricted to being presented by either a single allele in the case of single-allelic data, or one of the families of alleles (DR/DP/DQ). 10 negatives were randomly sampled from the proteome for each presented epitope in the mass spectrometry data. The negative samples were generated such that, for a given allele or set of alleles for multi-allelic samples, no sampled negative contained a nested positive epitope as a subsequence inside it. When validating EDGE-II trained using immunoaffinity capture data on the Reynisson *et al.* hold-out validation set^11^, we removed any peptide in the latter that shared a nonamer with any peptide in the same allele.

### EDGE-II mass spectrometry training data

#### Tumor and cell line specimens for EDGE-II training

Frozen tissue specimens for mass spectrometry analysis were obtained from commercial sources or hospital centers. A subset of samples collected were approved through Partners Healthcare IRB or under a research protocol approved by the Comité de Protection des Personnes, Ile-de-France VII. Written informed consent was obtained from all clinical study subjects whose biological samples were used in this study, and tissue from biobanks was obtained in compliance with all applicable laws. EBV transformed B cell lines expressing class II alleles were obtained from La Jolla Institute for Allergy and Immunology (La Jolla, CA).

### Immunoaffinity capture and isolation of class II peptides

Frozen tumors (0.1 mg – 1g) were prepared for analysis by first pulverizing liquid nitrogen snap-frozen tissues with a CP02 cryoPREP automated dry pulverizer (Covaris, Woburn, MA). Approximately 5% of the powdered tissue sample was placed in a vial for Smart-seq sequencing (performed as described in manufacturer’s protocol: Low Input RNA: cDNA Synthesis, Amplification and Library Generation (NEB #E6420)) and the remainder was used for immunoprecipitation (IP) enrichment of HLA peptides. Frozen cell pellets (0.5×10^8^ – 5×10^8^ cells) or pulverized tissue for IP enrichment were homogenized in 5 mLs of lysis buffer (20 mM Tris, 150 mM sodium chloride, 1% CHAPS, pH8) using a hand-held homogenizer (15-340-167, Thermo Fisher Scientific, Waltham, MA) and then rotated for 1 h to lyse. Lysates were spun at 21,000 x *g* for 60 min to clear cell debris, then added to pre-equilibrated filter plates containing NHS-sepharose beads coated in antibodies L243 (pan anti-HLA-DR, 94069, Biolegend, San Diego, CA), B721.1 (pan anti-HLA-DP) or SVPL3.1 (pan anti-HLA-DQ)). Both B721.1 and SVPL3.1 monoclonal antibodies were generated in SCID mice and provided by La Jolla Institute for Allergy and Immunology (La Jolla, CA) using their banked and validated hybridoma lines. Lysate was added and allowed to drip through a stack of plates prepared with each antibody using gravity flow. Then, plates were separated, and beads were washed twice with 1.8 mL 20 mM Tris, 150 mM NaCl, twice with 1.8 mL 20 mM Tris, 500 mM NaCl, and twice with 1.8 mL 20 mM Tris. Peptides were eluted with 200 µL of 2 N Acetic acid, then were purified using Sep-Pak tC18 plate (Waters Corporation, Milford, MA, 186002319) followed by an Oasis PRiME HLB µelution plate (Waters Corporation, 186008052) using 30% acetonitrile in 0.1% formic acid elution according to manufacturer’s guidelines. The peptides were dried down and stored at - 20°C prior to use.

### Mass spectrometry analysis of EDGE-II training samples

Dried class II immunopeptides were resuspended in 10 µL of 3% acetonitrile, 0.1% formic acid. A total of 4 µL of each sample was analyzed by mass spectrometry using an EASY-nLC 1200 (Thermo Fisher Scientific, Waltham, MA) coupled to an Orbitrap Fusion Lumos Tribrid mass spectrometer (Thermo Fisher Scientific) with a Nanospray Flex ion source (Thermo Fisher Scientific). The EASY-nLC was setup with a 75 µm x 2 cm precolumn (Thermo Fisher Scientific, 164946) followed by a 75 µm x 25 cm C18 analytical column (Thermo Fisher Scientific, 164941). A 180 min linear gradient starting with 4% acetonitrile, 0.1% formic acid and ending at 40% acetonitrile, 0.1% formic acid was used to elute the sample. Data was acquired in the Orbitrap at 120,000 resolution using full scan mass/charge (m/z) range of 350-2000 and 4E5 AGC target, followed by data dependent fragmentation HCD scans at 15,000 resolution, AGC target of 5E4, and collision energy 28 for charges 2-6. For the latter third of MS files collected, a data dependent decision tree was implemented which included electron transfer dissociation (ETD) MS2 scans in addition to HCD scans-based on charge. Briefly, electron transfer/higher energy (EThcD) fragmentation set at 15,000 resolution and 5E4 AGC, was triggered in separate m/z precursor scan ranges forz=3, 4, and 5-6.

### EDGE-II model

#### Large protein language model

EDGE-II leverages the pretrained protein sequence embeddings from the “Evolutionary Scale Model 2” (ESM2) language model^33^. ESM2 is trained using masked language modeling, which models the conditional distribution of a masked position inside a protein sequence, given the remainder of the sequence. The model was trained to minimize the negative log-likelihood of each true amino acid x_i_ given the remaining unmasked sequence *x*_\*M*_^31^.

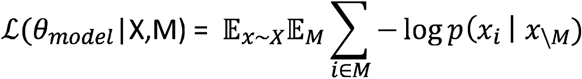

### Training

EDGE-II was implemented in Pytorch (v1.12.1) and consists of three subnetworks arranged in a feedforward structure: a sequence embedding network, a learned allele network (LAN), and a classification multilayer perceptron (MLP). The sequence embedding network, *f_θ_*, was initialized with the pre-trained weights *θ_ESM2_* from the ESM2 large protein language model ^33^. The input into the embedding network is an epitope sequence concatenated with a set HLA pseudosequences^11,48^. In the single allele case, a single epitope:allele combination is passed through the network. In the multi-allelic case, a given epitope is concatenated with each allele pseudosequence from the set of *K* alleles separately.

MA inputs were evaluated using a multiple instance learning (MIL) framework. Each MA sample, *x* ∊ *χ*, consisted of a set of peptide:alleles in a sample’s genotype, 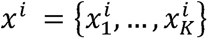, with a corresponding label *y^i^* ∊ {0,1} forming a training set ***B*** = {(*x*^1^,*y*^1^),…,(*x^N^,y^N^*)}. To train the network, MIL frameworks take the following general form to calculate a score s for which backpropagation against the loss function can be performed:

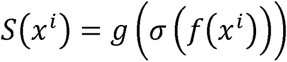

In the case of maximal-score deconvolution, *f* represents the ESM2-MLP classification network that produces *y* ∊ {0,1}. For EDGE-II with the LAN, *f* is the sequence embedding network (ESM2) and produces an intermediate representation *f*(*x^i^*) = *z^i^* ∊ ℝ^*d×n*^, where *d* is the embedding dimensionality of ESM2 and *n* is the number of alleles in the genotype. (J is any permutation invariant pooling operator that reduces the input set ℝ^*d×n*^ ➔ ℝ^*d*^, where *d* = 1 for maximal-score deconvolution and *d* = 480 for the LAN. σ is the max function for maximal-score deconvolution and for the LAN:

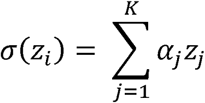

Finally, for maximal-score deconvolution *g* is simply the identity function and for the LAN *g* is the classification MLP. In the single allele case, the LAN reduces to the identity function. In both the SA and MA cases, a special classification token ([CLS]) was prepended to the start of each input was used as the final sequence representation.

The model was trained by iteratively exposing EDGE-II to three decreasing levels of presenting allele resolution. The first stage of training used exclusively single allelic data from the Reynisson *et al.* dataset^34^, followed by DR/DQ/DP-specific immunoaffinity purified mass spectrometry data (Methods), and the final round of training was on the Reynisson *et al.* dataset^34^. The Reynisson *et al.* MA dataset is primarily composed of full class II genotypes, necessitating the model to fully deconvolve up to 12 alleles. The LAN was included in this final round of training, prior to this step maximal score deconvolution was used on the DR/DP/DQ-specific data. All training was performed using the AdamW^49^ optimization algorithm with a learning rate of 0.0001 and weight decay of 0.001. Early stopping was used with validation average precision (AP) as the stopping criterion with a patience of 3 epochs. The best performing model was recovered after early stopping.

Confidence intervals for model performance metrics were calculated in Julia using bootstrap confidence intervals, with 1,000 resamples of the data.

### Predictiveness of EDGE-II scores with immunogenicity

Bayesian explanatory logistic regression was used to measure the association between EDGE-II, NetMHCIIpan4.3^19^, MixMHC2pred-2.0^20^, and CD4 immunogenicity data from a previously published individualized cancer vaccine study^35^. 50% and 90% credible intervals for the regression weight between presentation score and immunogenicity label were calculated by sampling the posterior distribution of the trained regression model.

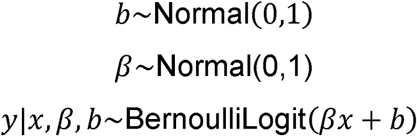

The model was implemented in and MCMC sampling performed using the Julia based Turing.jl probabilistic programming language^30,50,51^. The NUTS sampling algorithm was used with mean acceptance probability of 0.65 (default), with 2,000 samples drawn after a 1,000 sample burn-in period. Normal(0,1) priors were used for the parameters (coefficient and intercept) in the logistic regression. Due to the difference in scale of scores reported by EDGE-II (logits), NetMHCIIpan4.3 (probability and rank percentile), and MixMHC2pred-2.0 (rank percentile), input scores from the models were standardized by subtracting the mean and dividing by the standard deviation for each. For plotting purposes, the standardized scores were converted back to their original values.

### Gaussian process to visualize EDGE-II scores as a continuous function of HLA sequence

Embeddings for each HLA pseudosequence were extracted from ESM2 using mean-pooling over the length dimension. UMAP was then performed on these embeddings and plotted in 2D. A Gaussian process (GP) was fit to EDGE-II scores using the UMAP allele embeddings as predictors to visualize EDGE-II scores as a function of HLA sequence. The Julia GaussianProcesses.jl (v0.12.5) library used with the MeanZero() mean function (no prior belief in the function expectation) and SquaredExponential (Gaussian) kernel function with log-scale parameters (1.0,-0.5).

### Saliency Mapping of KRAS G12C Epitopes in Patient S2

Positional importance for HLA class II presentation was estimated in Patient S2 using the method of saliency^52^, as implemented in the captum library (v0.5.0). All G12C containing 15-mers were conditioned against Patient S2’s HLA genotype and passed through EDGE-II. The gradient of the ESM2 embeddings was extracted for each peptide, averaged over all alleles, and summed over the ESM2 latent embedding dimension. This operation yields a scalar value at each position that was then multiplied by the presentation probability of peptide and normalized against the number of occurrences of each position in the set of G12C containing peptides.

### Human T cell data

#### Ethics statement

The research presented in this manuscript complies with all the relevant ethical regulations. The clinical study and all related analyses were carried out in accordance with the Declaration of Helsinki and Good Clinical Practice guidelines and was approved by the appropriate IRB or ethics committees at each Sarah Cannon Research Institute. All patients provided written, informed consent. Further details can be found at https://clinicaltrials.gov/ct2/show/NCT03953235.

### PBMC isolation and storage

Whole blood samples from a cancer patient were collected with informed consent, IRB approval, and according to protocol during an investigational Phase 1/2 dose-finding study (<u>NCT03953235</u>)^28^. PBMCs from whole blood were isolated isolated using density gradient centrifugation on Ficoll® Paque Plus (GE Healthcare, Chicago, IL, USA), washed with D-PBS (Corning, Corning, NY, USA), counted, and cryopreserved in CryoStor CS10 (STEMCELL^TM^ Technologies) at 5 x 10^6^ cells/ml. Cryopreserved cells were stored in liquid N_2_ (LN_2_), shipped in cryoports and transferred to storage in LN_2_ upon arrival. Cryopreserved cells were thawed and washed twice in OpTmizer^TM^ T Cell Expansion Basal Medium (Gibco, Gaithersburg, MD, USA) with Benzonase (EMD Millipore, Billerica, MA, USA) and once without Benzonase. Cell counts and viability were assessed using the Guava ViaCount reagents and module on the Guava EasyCyte HT cytometer (EMD Millipore). Cells were rested overnight prior to use in functional assays.

### HLA typing (class I and II)

HLA class I typing was performed on patient samples as part of enrollment in the GO-005 clinical trial. Whole blood samples were collected at the patient’s collection site and processed for cell free DNA (cfDNA). CfDNA was sent to University of California – Los Angeles (UCLA) for whole gene HLA typing utilizing NGS-based HLA typing methods to achieve high resolution class I and class II full gene sequencing. Only class I results were reported to the Sponsor.

HLA class II typing was performed in-house. HLA typing protocol was completed using kits by GendX (HLA-DRB3/4/5: Cat. 7370762, HLA-DPA1: Cat. 7370962, HLA-DQA1: Cat. 7370862, NGSgo-LibrX Cat. 2343605) and following manufacturer’s established protocol (Amplification protocol: IFU NGSgo-AmpX v2 7310000 edition 7, 2022-07 RUO. Library preparation protocol: IFU NGSgo Library Full Kit 2312006 edition 3, 2022-06 RUO). A subset of samples were typed from bulk RNAseq using arcasHLA (https://academic.oup.com/bioinformatics/article/36/1/33/5512361) v0.1.1, discarding DRB3/4/5 types which did not agree with the DRB1 types.

### Peptides

Custom-made, recombinant, lyophilized peptides specific for G12C mutation were produced by Genscript (Piscataway, NJ, USA) and reconstituted at 2 or 5mg/ml/peptide in sterile DMSO (VWR International, Pittsburgh, PA, USA), aliquoted, and stored at - 80°C. Control peptides to assess responses to infectious disease antigens from CMV, EBV, Influenza (CEF peptide pool) were purchased from JPT Peptide Technologies (Berlin, Germany).

### Depletion CD4+ or CD8+ T cells from PBMCs

PBMC samples resuspended at 2×10^6^ cells/mL were divided into three different tubes to allow for three conditions: untouched (total) PBMCs, CD4^+^ depleted, and CD8^+^ depleted PBMCs. PBMCs set aside for the depletion process were centrifuged at 300 x g for 10 minutes at room temperature. Samples were resuspended in 80uL of MACS Buffer (1:20 dilution of MACS Rinsing BSA Stock Solution ‘CAS: 130-091-376’ into autoMACS Rinsing Solution ‘CAS: 130-091-222’) per 10^7^ total cells. PBMCs were labeled for depletion by adding 20µL of either CD4 human MicroBeads (CAS: 130-045-101) (CD4^+^ depletion) or 20µL of CD8 human MicroBeads (CAS: 130-045-201) (CD8+ depletion) per 10^7^ PBMCs. Cells were mixed with MicroBeads and placed at 4°C for 15 minutes. After incubation, 1-2 mL of MACS buffer was added to each conical and cells were spun at 300 x g for 10 minutes at room temperature. Each sample was resuspended in 500µL of MACS buffer (per 10^8^ PBMCs) prior to magnetic separation. The following steps were completed using Miltenyi Biotec MS columns (CAS: 130-042-201), Pre-Separation Filters (CAS:130-041-407), and OctoMACS separation magnet (CAS: 130-042-109). MS Column with Pre-Separation filter was placed on top of MS columns inserted into a Miltenyi magnet. Filters and columns were washed with 500µL of MACS buffer, allowing for the buffer to completely drip and soaking the column and filter. Each labeled cell suspension was added to a separate column and flow-through was collected. Once column reservoir was empty, columns were washed 3 times with 500uL MACS buffer and effluent was collected in the same collection tube. After washes were complete, cells were resuspended in assay media and stimulated as described below. Depleted cell suspensions and untouched PBMC samples were phenotyped to ensure complete depletion of unwanted cell population using anti-CD4^+^ (APC-ef780 Ref: 47004942), anti-CD8^+^ (Percp-Cy5.5 Ref: 300924), Live/Dead (Zombie Red Ref: I23101), and anti-CD3^+^ (BV421 Ref: 300434) antibodies. Samples were acquired on the Cytoflex flow cytometer and analyzed using FlowJo analysis software.

### *In vitro* stimulation (IVS) cultures

Neoantigen-reactive T cells from patient samples were expanded in the presence of cognate peptides and low-dose IL-2 as described previously^3^. Briefly, thawed PBMCs were rested overnight and stimulated in the presence of minimal epitope peptide pools (10µg/ml/peptide), class II peptides (4ug/ml/peptide) or control peptides (CEF) in ImmunoCult™-XF T Cell Expansion Medium (IC media; STEMCELL Technologies) with 10 IU/ml rhIL-2 (R&D Systems Inc., Minneapolis, MN, USA) +/- 0.5 µM bME for 14 days in 48- or 24-well tissue culture plates. Cells were seeded at 1-2 x 10^6^ cells/well and fed every 2-3 days by replacing 2/3 of the culture media with fresh media containing rhIL-2.

### IFNg ELISpot assay

Detection of IFNg-producing T cells was performed by *ex vivo* ELISpot assay. Briefly, cells were harvested, counted and re-suspended in media at 4 x 10^6^ cells/ml and cultured in the presence of DMSO (VWR International), Phytohemagglutinin-L (PHA-L; Sigma-Aldrich, Natick, MA, USA), CEF peptide pool, or cognate peptides in ELISpot Multiscreen plates (EMD Millipore) coated with anti-human IFNg capture antibody (Mabtech, Cincinnati, OH, USA). Following 18-24h incubation in a 5% CO_2_, 37°C, humidified incubator, supernatants were collected, cells were removed from the plate, and membrane-bound IFNg was detected using anti-human IFNg detection antibody (Mabtech), Vectastain Avidin peroxidase complex (Vector Labs, Burlingame, CA, USA) and AEC Substrate (BD Biosciences, San Jose, CA, USA). Plates were imaged and enumerated on an AID iSpot reader (Autoimmun Diagnostika GmbH, Strassberg, Germany). Data are presented as spot forming units (SFU) per million cells.

### Flow cytometry analyses

Samples acquired on flow cytometers were analyzed using FlowJo software (FlowJo, LLC, Ashland, OR, USA). Gating strategies for human samples and Jurkat cell line TCR functional screening assays are as follows: **CD4 versus CD8 T cell assessments**: Lymphocytes (SSC-A vs FSC-A), single cells (FSC-H vs FSC-A), viable cells (FSC-A vs LD-ZombieRed), CD3^+^ cells (FSC-A vs CD3-BV605), CD4^+^ and CD8^+^ (CD4-APCeF780 vs CD8-PerCP-Cy5.5). **ICS studies**: Lymphocytes (SSC-A vs FSC-A), single cells (FSC-H vs FSC-A), viable cells (FSC-A vs LD-ZombieRed), CD3^+^ cells (FSC-A vs CD3-BV605), CD4^+^ and CD8^+^ (CD4-APCeF780 vs CD8-PerCP-Cy5.5), CD8^+^ or CD4^+^ cytokine^+^ cells (CD8-PerCP-Cy5.5 vs CD107a-BV421, IFNg-APC, TNFa-BV786, IL-2-PE; CD4-APCeF780 vs CD107a-BV421, IFNg-APC, TNFa-BV786, IL-2-PE).

### Single cell TCR sequencing and digital gene expression

Single-cell library preparation and sequencing was completed according to 10x Genomics v1 5’ Chromium Single Cell V(D)J protocol CG000186 Rev A (Pleasanton, CA, USA). The only deviation in library preparation was in V(D)J library step 5.4 with 5 cycles of PCR. All sequencing was completed on the Illumina NovaSeq at a loading concentration of 450 pM with 5% PhiX (San Diego, CA, USA). The 10x Genomics Cell Ranger version 3.1 was used for V(D)J and Digital Gene Expression library analysis and was current at time of research.

### Single cell RNAseq data analysis

The single cell RNAseq data was processed using 10x Genomics Cell Ranger v3.0.2^53^. Using the Cell Ranger count pipeline and human reference (GRCh38, cellranger reference 1.2.0, Ensembl v84 gene annotation) the feature barcode expression matrix was generated for each sample. R (v4.2.0; https://www.R-project.org/) based Seurat (v4.1.1)^54^. pipeline was used to filter low quality cells, filter non-T cells, cluster and annotate T cells. First, Seurat Read10x() function was used to read the cell ranger output files. Next, the expression matrices representing an individual patient were merged into one large expression matrix. The CreateSeuratObject() function was used to create a Seurat object by loading the patient specific expression matrix. To filter out the low-quality cells and non-T cells a series of clustering and filtering steps were performed. In order to delete the dead cells and low quality cells, the standard Seurat clustering steps (https://satijalab.org/seurat/articles/pbmc3k_tutorial.html; Date: May 21, 2022) were performed using 20 PCAs and 0.8 resolutions to generate cell clusters. Dead cell clusters were identified by measuring the proportion of mitochondrial gene expression in each cluster. If the proportion of mitochondrial gene expression in a cluster was three times the average mitochondrial genes expression proportion in all the cells, and eight out of top ten differentially expressed genes in that cluster were mitochondrial genes, we define that cluster as a dead cell cluster. All the cells in the dead cell cluster were filtered out. In addition, low quality cells were filtered out using the following parameters in subset() function (nFeature_RNA > 600; nFeature_RNA < 6000; nCount_RNA < 40000; percent mitochondrial genes < 20; percent ribosomal genes > 5). In order to delete the non-T cell, we use SCTransform()^55^ function to normalize gene expression and regressed the influence of technical characteristics such as, Cell Cycle scores (S.score and G2M.score), expression percentages of mitochondrial and ribosomal genes. Next, batch correction was performed using RunHarmony()^56^ function. Single cell clusters were generated using 25 PCAs and 0.8 resolutions. From the resulting clusters we identified the differentially expressed genes (min.pct = 0.25, logfc.threshold = 0.25) in each cluster. The R package CIPR (v0.1.0)^57^ was used to annotate the cell type of each cluster, the differentially expressed genes in each cluster were compared to the differentially expressed genes in one of the reference datasets defined within CIPR (comp_method = "logfc_dot_product", reference = "hsrnaseq") to define cell types that best represented the given clusters. The clusters annotated as non-T cells were filtered out. Only the cells which had at least one read representing CD3D, CD3E, CD8A, CD8B or CD4 genes were defined as T cells and considered for downstream analysis. With the low quality and non-T cells filtered, the T cells were normalized using SCTransform()^55^ function and the influence of technical characteristics such as, Cell Cycle scores (S.score and G2M.score), nCount_RNA, expression percentages of mitochondrial and ribosomal genes were regressed out. The variable features excluding the TCR receptor genes TR[ABDG]V were used to obtain PCAs. The experimental variables like sample name, dose (D1, D2, etc), IVS(CEF, PEPTIDE), ELIS (stimulated, DMSO) were considered during batch correction process using RunHarmony(). The single cell clusters were generated using 35 PCAs and 0.8 resolutions. RunUMAP() was used for UMAP dimensionality reduction. From the resulting clusters, differentially expressed genes (min.pct = 0.25, logfc.threshold = 0.25) were identified for each cluster. The cell type annotation was performed through combination of cell type markers, CIPR results and literature review of differentially expressed genes in each cluster. FeaturePlot(), DimPlot(), DoHeatmap(), and VlnPlot() functions from Seurat were used to generate different visualizations.

### Single cell TCRseq data analysis

The single cell TCR sequencing data was processed using 10x Genomics Cell Ranger v3.0.2^53^. Using the Cell Ranger vdj pipeline and V(D)J reference (Human GRCh38 V(D)J sequences, cellranger reference 2.0.0, annotation built from Ensembl Homo_sapiens.GRCh38.87.chr_patch_hapl_scaff.gtf and vdj_GRCh38_alts_ensembl_10x_genes-2.0.0.gtf) the TCR clonotypes were generated for each sample. R package scRepertoire (v1.7.2)^58^ was used to process the TCR output files from Cell Ranger and integrated the TCR information into the Seurat object at single cell resolution. The combineTCR() function was used to select the T cells with alpha and beta chain sequences and filter out T cells with missing values or gamma/delta T cells. TCR annotations of samples obtained from a single patient were combined into one patient specific TCR annotation matrix for downstream integration into single cell expression data.

### Integration of single cell TCR and single cell gene expression data

The Seurat object at the end of single cell RNAseq analysis and the TCR annotation from the single cell TCRseq analysis were merged using the combineExpression() function in scRepertoire. The parameter cloneCall="aa" was used which considers the combination of amino acid sequence of alpha and beta chain CDR3 sequences as the unique id for each TCR sequence, and the frequency of TCR clones were obtained. The function also bins each T cells with TCR sequence annotation into sub-clonotypes based on their frequency in a given sample: Hyperexpanded (100 < X <= 500)", "Large (20 < X <= 100)", "Medium (5 < X <= 20)", "Small (1 < X <= 5)", "Single (0 < X <= 1)", "NA".

### Cell lines

The cell lines K562 are lymphoblastic cells isolated from the bone marrow of a myelogenous leukemia female patient. K562 were maintained on Iscove’s Modified Dulbecco’s Medium (IMDM) supplemented with 10 % fetal bovine serum (FBS gibco: Lot#2364822P) plus 1 % of Penicillin/streptomycin 50 mg/mL (Gibco: 10378016). The KAS116 is a lymphoblastoid cell line isolated from the blood of a human female patient. The Jurkat TCRKO Nur77-GFP is transgenic version derived from Jurkat clone E.1, which was obtained from a young male with acute T cell leukemia. KAS116 cell lines and the transgenic Jurkat cell lines were maintained on RPMI-1640 medium supplemented with 2mM L-Glutamine, HEPES (fc 10mM), Penicillin/streptomycin (FC 50U/ml) and 10% of FBS. All the lines were culture at 37 °C and 5% of C_2_O. the initial culture at 1×10^5^ cells/ml in a T-75 flasks (Corning #431464) subculture at 1x 106 cell/ml every 2-3 days.

### Recombinant TCR cloning, virus production, and transductions

TCR sequences (From the 10x sequencing) were cloned into a pCD502_MSCV vector backbone containing the features for lentiviral transduction (TWIST Bioscience) using Gibson Assembly. Cloned vector was received in house and resuspended in nuclease free water (REF AM937). Plasmids were utilized to generate lentivirus in 96-well plates. Lenti-X™ 293T cells (Takara 632180) were utilized for transfection and virus production. 24 hours prior to transfection, 30,000/well Lenti-X 293T cells were plated in a 96-well flat bottom plate (REF 3548). Transgene plasmids (25ng) were added to 10 μL of OptiMEM (Gibco 31985062) along with 60ng of the packaging plasmid ViraPower ™ Lentiviral Packaging Mix (Invitrogen K497500). Separately, 0.25 μL of Lipofectamine 2000 (Invitrogen 11668019) to 10 μL of OptiMEM. These were incubated separately for 5 minutes, combined, and then incubated for another 20 minutes together. Transfection reagents were then added slowly drop-wise to the Lenti-X 293T cells previously plated 24 hours prior. Viral supernatant was left undisturbed for 48 hours. After 48 hours of incubation, supernatant was harvested and filtered through a 0.45 μm PES filter to remove any 293T cells. A Jurkat E6.1 (ATCC TIB-152) derived cell line was generated by CRISPR knocking out of endogenous TCR alpha and beta genes. This cell line was used for CD4 + TCR studies. For utilizing CD8 + TCRs, CD8 isoform 2 was transduced with lentivirus and CD4 was knocked out using CRISPR to generate solely CD8 + T cells. 30,000 of CD4 or CD8 Jurkat cells were plated into round bottom 96-well plates along with 8μg/mL of Polybrene (EMD Millipore Cat # TR-1003). Cells were spinoculated at 800 x g for 45 minutes. Cells were grown for 72 hours and then checked for TCR expressing using APC anti-human CD3 (BioLegend Clone UCHT3 Cat# 300439).

### Recombinant TCR screening assays (Jurkats)

Jurkats with recombinant TCRs were prepared as shown above in “Recombinant TCR Cloning, Virus Production, and Transduction.” Target cells utilized in class II TCR screening is a DRB1*01:01 HLA matched B cell line, Kas116 (Millipore Sigma 88052003-1VL; Obtained from Alexander Sette’s lab). Target cells (Kas116) were counted and plated into a flat bottom 96-well plate at 75,000 cells/well in 100 μL of OpTimizer™ (Gibco A1048501). 20 μM of test peptide was also added to appropriate wells +/- 0.05 µM bME. The plate was then placed into an incubator to pulse peptide for approximately 1 hour. After the hour of peptide pulse, add 75,000 cells of Jurkat cells to each corresponding well for a 1:1 Effector to Target Ratio (E:T). Incubate together for 20 hours overnight. Post incubation, collect the wells and transfer to a round bottom 96-well plate to begin staining. Spin the plate at 500 x g for 3 minutes. Flick off supernatant and resuspend in 100 μL of EasySep buffer (StemCell 20144) to wash. Spin again at 500 x g for 3 minutes. Flick off supernatant again. Resuspend in 1:100 diluted antibody solution containing the following: PE anti-human CD25 (BioLegend Clone BC96 Cat# 302606), APC anti-human CD69 (BioLegend Clone FN50 Cat# 310910) and then either: BV421 anti-human CD8a (BioLegend Clone RPA-T8 Cat# 301036) or BV421 anti-human CD4 (BioLegend Clone RPA-T4 Cat# 300532). Stain at room temperature for 15 minutes in the dark. Add 100 μL of EasySep buffer to wash and spin at 500 x g for 3 minutes. Flick off and then resuspend in 200 μL of EasySep buffer and proceed to acquire flow. Polyfunctionality analyses were performed using Boolean gating (FlowJo). Results are represented as % positive cell populations (frequency of parent). Data shown as background subtracted where indicated. Gate on Lymphocytes (FSC-A vs SSC-A), Single Cells (FSC-A vs FSC-H),, CD69 (APC-A vs FSC-H), CD25 (PE-A vs FSC-H) and Nur-77 GFP (FITC-A vs FSC-H)

### Data visualizations

Figures 1,2, and 6AE were visualized using Julia (Plots.jl,StatsPlots.jl). The remaining graphical representations of data were performed using GraphPad Prism 9 for macOS (Version 9.0.1 (128). Schematics were created with <u>BioRender.com</u>. Representative flow cytometry dot plots and polyfunctionality pie charts were graphed using FlowJo Version 10.6.1 for Mac OS X.

## Acknowledgements

Fatema Z Chowdhury, Aaron Yang, Raj Khatri, Charuta Yadav, Minh Duc Cao, Charmaine N Nganje, Aleksandra Nowicka, Elizabeth Maloney, Martina Marrali, Harshni Venkatraman, Jason R Jaroslavsky, Cassidy Jones, Aarushi Bhan, Tyler Murphy, Lindsey Arcebuche, Lorenzo Hernandez, patient & family, clinical staff at SCRI, Kamilah Caldwell & Cynthia Voong, Alison Dixon.

## Competing Interests

D.S, M.G.H., J. K., S. K., R. V., M. R., L. D. K.T., R.Z., L. K., A. C. G., J. A., A. M., B. A., C. D., A. R. F., M. L. J., M. J. D., M. L., C. D. P., K. J., and A. D. are stockholders and either current or previous employees at Gritstone bio, Inc. and may be listed as co-inventors on various pending patent applications related to the methods presented in this study. M.L.J.: financial interests, personal, advisory board: Astellas, Otsuka; financial interests, institutional, research grant: AbbVie, Acerta, Amgen, Apexigen, Arcus, Array, AstraZeneca, Atreca, Beigene, Birdie, Boehringer Ingelheim, Checkpoint Therapeutics, Guardant Health, Genocea, Hengrui, Immunocore, Incyte, Janseen, Jounce, Gritstone bio, Lycera, Merck, Mirati, Oncomed, Regeneron, Ribon, Sanofi, Shattuck Labs, Stem CentRx, Syndax, Takeda, Tarveda, TCR2 Therapeutics, University of Michigan and WindMIL.

## Supplementary Figures

**Supplementary Figure S1:**
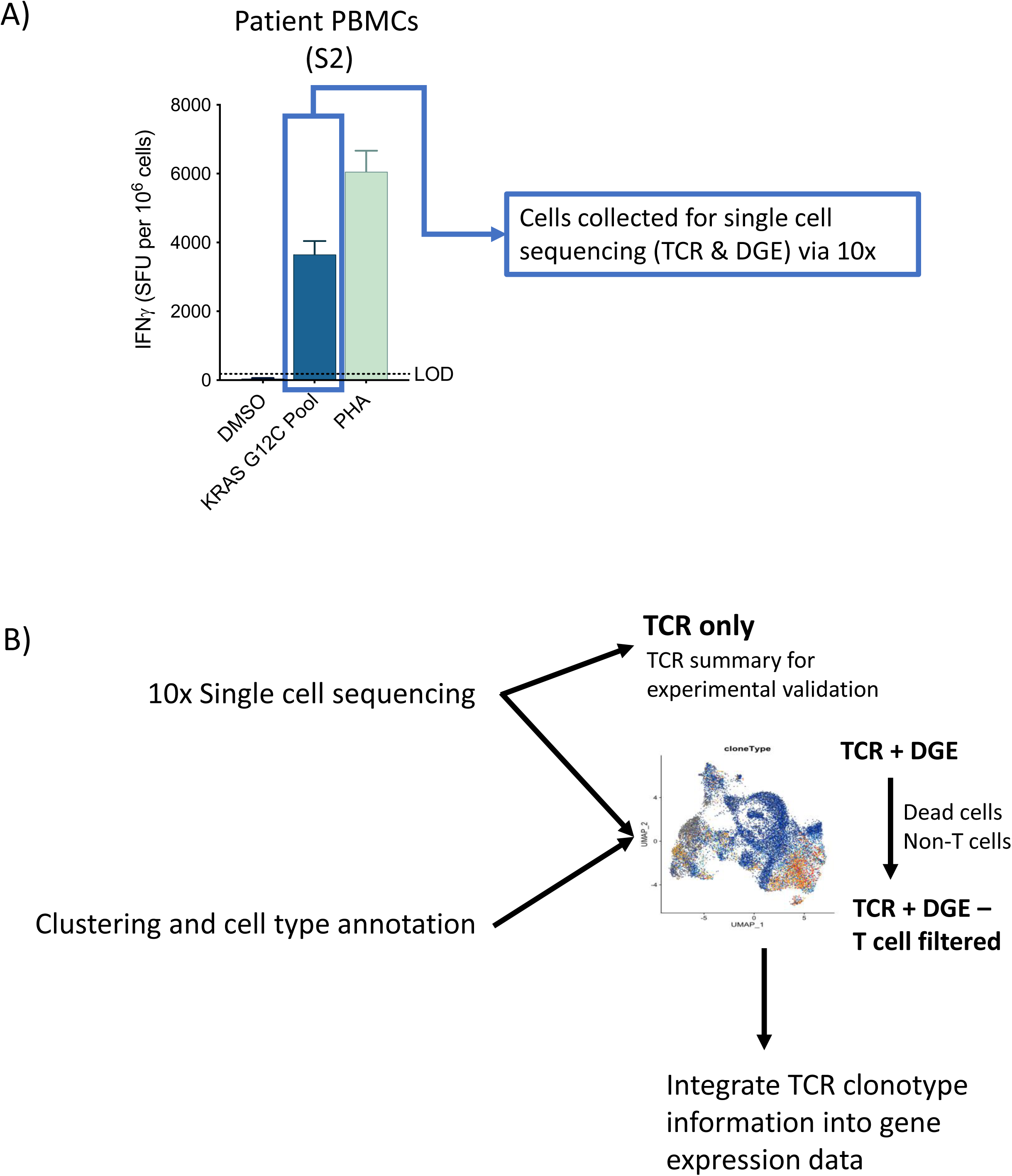
Sample collection and analysis for Patient S2 TCRseq and Digital Gene Expression (DGE) analysis. **A)** Post-immunotherapy T cell responses to KRAS G12C peptide pool (Supplementary Table S5) and controls assessed by post-IVS IFNγ ELISpot for Patient S2. Bar graphs (mean ± SD of replicate wells) show SFU/10^6^ cells. Assay limit of detection (LOD; 180 SFU/10^6^ cells) is indicated by dotted line. Box and arrow indicate post-ELISpot cells collected for 10x emulsions and single cell sequencing. B) Schematic outlining single cell sequencing approach for TCR seq and digital gene expression (DGE) analyses.

## Notes

### Summary of Updates

Figure 1 revised, Figure 2 revised

